# Exploring Okra-Derived Compounds as Prospective Aromatase Inhibitors: A Computational Study for Enhanced Breast Cancer Therapy

**DOI:** 10.1101/2023.10.25.563890

**Authors:** B Lavanya, Dhrithi Jayasimha Mallur, Sheshadri S Temkar, V Arun, Benedict Paul C

## Abstract

Breast cancer with estrogen receptor positivity represents itself as the most prevalent malignancy among postmenopausal women. One of the promising therapeutic approach involves the use of Aromatase Inhibitors, which competitively bind to aromatase, reducing estrone and estradiol levels. While current drugs have improved survival rates, they are not without adverse effects. Consequently, this study explores the computational screening of medicinally relevant compounds derived from Okra for potential Aromatase Inhibition. Molecular Docking, employing AMDock v1.5.2, was utilized to assess binding affinities with aromatase (PDB:3EQM). Subsequently, in-depth molecular interactions were examined using Discovery Studio Visualizer v4.5, and the stability of docked complexes was evaluated via Molecular Dynamics with the GROMACS package, focusing on RMSD, RMSF, H-bond count, and SASA. The pharmacokinetic properties of the Okra compounds were predicted using admetSAR v2.0. Our findings highlight Quercetin 3-gentiobioside as a standout candidate, demonstrating superior binding affinity (-10 kcal/mol) and an Estimated Ki of 46.77 nM compared to Letrozole and other Okra compounds. Molecular dynamics analysis confirms the stability of Quercetin 3-gentiobioside binding in terms of H-bonds and conformational integrity. In conclusion, our computational investigation identifies Quercetin 3-gentiobioside, along with Quercetin 3-O-rutinoside and Hyperin, as promising candidates for preclinical studies in the pursuit of potential Aromatase Inhibitors.

## Introduction

Breast cancer with estrogen receptor (ER) positivity is the predominant malignancy among postmenopausal women and is associated with a higher mortality rate compared to ER-negative cases, even in non-malignant conditions. Current therapeutic strategies for ER-positive breast cancer include selective estrogen receptor modulators (SERMs) that antagonize estrogen action, selective estrogen receptor degraders (SERDs), and aromatase inhibitors (AIs) that inhibit estrogen synthesis (Haynes *et al*., 2003). AIs, such as anastrozole and letrozole, have emerged as powerful third-generation options, serving as primary therapies in recent ER-positive breast cancer cases with relatively tolerable side effects compared to earlier treatments. Nevertheless, AIs are not without adverse effects, which include hot flashes, vaginal dryness, osteoporosis, menopausal symptoms, musculoskeletal discomfort, and an elevated risk of bone fractures as long-term consequences (Mortimer, 2010; Serban *et al*., 2022). Some cases also report ocular and neuro-retinal impairment (Ağın *et al*., 2021). Therefore, the discovery of novel AIs is crucial for improving the management of hormone-sensitive breast cancer.

Studies indicate that plant-derived compounds, often used in combination with conventional anticancer drugs, hold promise in selectively targeting tumor cells while sparing normal cells, such as lymphocytes and fibroblasts (Lichota and Gwozdzinski, 2018). Some research even suggests that certain plant extracts possess potent standalone anticancer properties (Kamatou, 2008; Ochwang’i, 2014; Naik and Kumar, 2020). Consequently, plant phytocompounds offer a valuable resource for alternative therapeutic strategies, potentially mitigating the side effects associated with traditional drug treatments.

Many chemotherapeutic drugs used in cancer treatment are synthetic derivatives of various plant compounds, including *Vinca alkaloids*, *Taxus diterpenes*, and *Camptotheca alkaloids*. These plant compounds inhibit tumor cell growth through selective mechanisms such as apoptosis induction, DNA damage, and inhibition of topoisomerase I and II (Lichota and Gwozdzinski, 2018). In addition to alkaloids and diterpenes, flavonoids are widely distributed in vascular plants and have garnered attention for their potential for disease risk reduction. These polyphenolic compounds possess various health-promoting effects, including antioxidative, anti-inflammatory, antimutagenic, and anticarcinogenic properties (Panche *et al*., 2016). Recent research has highlighted the role of flavonoids in cancer by modulating cell death signaling pathways through reactive oxygen species (ROS) regulation and apoptosis induction (Scarano *et al*., 2018; Abotaleb *et al*., 2018; Kopustinskiene *et al*., 2020). Among these flavonoids, quercetin stands out as a clinically relevant phytocompound with antioxidative and anticancer properties (Lesjak *et al*, 2018; Lamson and Brignall, 2000).

Okra (*Abelmoschus esculentus*), commonly known as Lady’s Finger, belongs to the Malvaceae family and is renowned for its nutritional value (Islam, 2019). Its green seed pods, consumed raw or cooked, are rich in protein and unsaturated fatty acids like linoleic acid. Okra also boasts a substantial amount of bioactive flavonoids, including quercetin and its derivatives. Among different plant parts, okra flowers exhibit the highest levels of total phenolics, total flavonoids, and antioxidant activities (Deng *et al*., 2020; Lin *et al*., 2014). Notably, flavonoid constituents from okra seed extracts demonstrate dose-dependent cytotoxic effects on MCF-7 breast cancer cells (Gemede *et al*., 2015).

With its abundance of bioactive compounds, okra presents itself as a promising source for health solutions. Compounds derived from okra have shown potential for reducing the risk of chronic diseases like atherosclerosis and cancer (Lichota and Gwozdzinski, 2018). Hyperin, found in okra, has been noted for its anticancer properties against gastric cancer cells (CHI), inducing apoptosis by blocking the Wnt/β-catenin signaling pathway (Elkhalifa *et al*., 2021).

The demand for highly effective and safe antitumor drugs is growing. Plant extracts and phytocompounds are increasingly explored for their medicinal and antitumor effects. In pursuit of non-toxic therapeutic solutions utilizing medicinally relevant plants, this study investigates okra phytocompounds as potential aromatase inhibitors. Therefore, we employ computational techniques to screen and assess the potential of okra phytocompounds in this regard.

## Methods

### Selection of Okra Compounds

A comprehensive literature review led to the identification of nine distinct compounds for investigation. The tertiary structures of these compounds were obtained from PubChem (Kim *et al*., 2019). The compounds, along with their respective PubChem IDs, are detailed in Table 1. Subsequently, these compounds were downloaded as SDF (Structure-Data File) formats and then converted to the PDB (Protein Data Bank) format using OpenBabel v. 3.1.1 (O’Boyle *et al*., 2011).

**Table 1:**
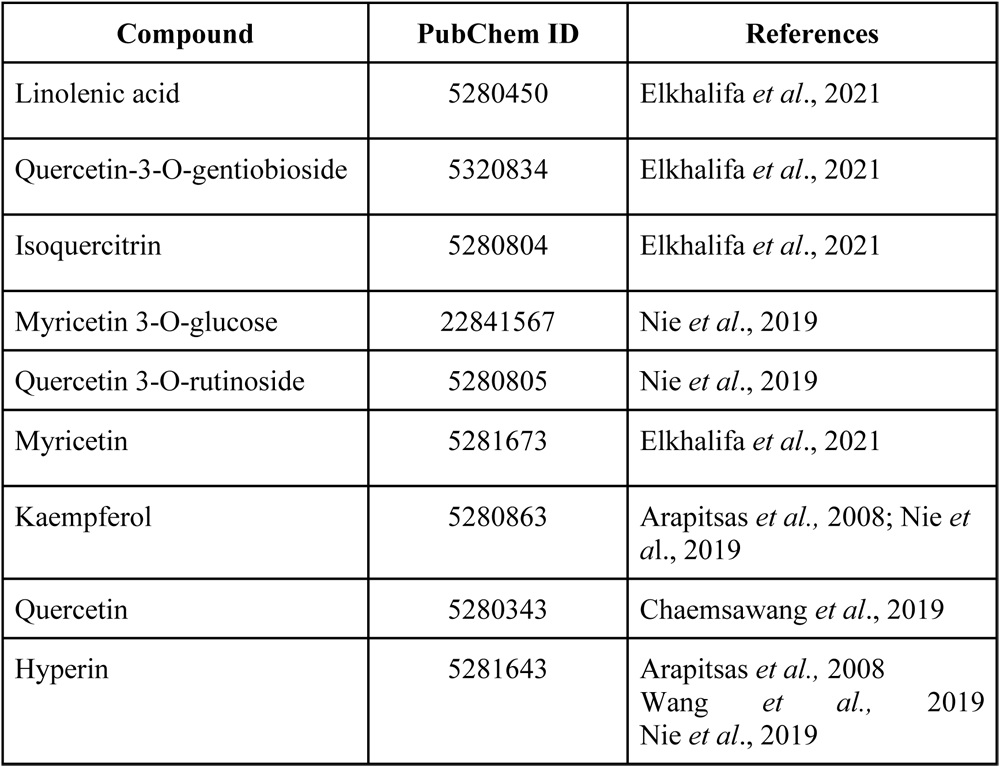
Selected Okra compounds for the study.

| Compound | PubChem ID | References |
| --- | --- | --- |
| Linolenic acid | 5280450 | Elkhalifa *et al.*, 2021 |
| Quercetin-3-O-gentiobioside | 5320834 | Elkhalifa *et al.*, 2021 |
| Isoquercitrin | 5280804 | Elkhalifa *et al.*, 2021 |
| Myricetin 3-O-glucose | 22841567 | Nie *et al.*, 2019 |
| Quercetin 3-O-rutinoside | 5280805 | Nie *et al.*, 2019 |
| Myricetin | 5281673 | Elkhalifa *et al.*, 2021 |
| Kaempferol | 5280863 | Arapitsas *et al.*, 2008; Nie *et al.*, 2019 |
| Quercetin | 5280343 | Chaemsawang *et al.*, 2019 |
| Hyperin | 5281643 | Arapitsas *et al.*, 2008<br>Wang *et al.*, 2019<br>Nie *et al.*, 2019 |

### Selection of Protein Structure from PDB

The three-dimensional crystal structure of human placental aromatase cytochrome P450 complexed with androstenedione (accession number: 3EQM) was retrieved from the Protein Data Bank (PDB) (Ghosh 2012, Berman *et al*., 2000).

### Docking Simulations of Native Protein with Okra Phytocompounds

Molecular docking studies were carried out using AMDock v.1.5.2, a versatile software that utilizes the AutoDock engine for molecular docking. AMDock incorporates essential tools such as Open Babel, PDB2PQR, and PyMol for structural analysis (Valdés-Tresanco *et al*., 2020). This tool was selected for its seamless docking capabilities and robust functionality. Docking results were evaluated in terms of binding affinity (kcal/mol), estimated Ki (µM), and ligand efficiency. The Ki values were calculated using the AutoDock Vina software with the equation exp(ΔG/RT), where R represents the universal gas constant (1.985 × 10^-3^ kcal mol^-1^K^-1^) and T is the temperature (298.15 K). The compound with the most negative binding affinity and the lowest Ki value was selected for each compound. The tertiary structure of the commercial nonsteroidal drug: Letrozole (PubChem CID: 3902) was used as a reference in all docking simulations.

### Elucidation of Molecular Interactions

The docked complexes were exported from AMDock in PDB format and analyzed for hydrogen and non-bonded interactions using Discovery Studio Visualizer v4.5.

### ADMET Analysis

Toxicity analysis of phytocompounds is crucial for their potential as drug candidates. Okra phytocompounds with the lowest estimated Ki values and the highest binding efficiency were subjected to ADMET analysis, which evaluated their toxicity and absorption levels. ADMET analysis was conducted using admetSAR (v2) (Cheng *et al*., 2012). The SDF files of the phytocompounds were obtained from PubChem and converted to their respective SMILES (Simplified Molecular Input Line Entry System) formats using OpenBabel. Factors such as oral toxicity, carcinogenicity, and blood-brain barrier penetration were considered in the ranking of phytocompounds. The oral toxicity of the compounds was assessed using the LD50 values based on the scale recommended by the United States Environmental Protection Agency. Toxicity levels were categorized as “Practically Non-Toxic,” “Slightly Toxic,” and “Toxic,” following the grading criteria suggested by the US Office of the Federal Register in 2012.

### Molecular Dynamics

Molecular dynamics (MD) simulations, a computational technique employing Newton’s laws of motion, were conducted to study the atom movements within the molecules. MD simulations were observed for (i) Protein [3EQM-APO], (ii) Protein bound to Androstenedione [3EQM-ASD], and (iii) Protein bound to our compound of interest, Quercetin-3-O-gentiobioside [3EQM-QUR]. The Gromacs software package, a well-established MD simulation software, was employed for this purpose (Van Der Spoel *et al*., 2005).

The simulation process began with the minimization of the protein-ligand complex in a vacuum using the steepest descent algorithm. This iterative process adjusted the atomic coordinates to minimize the system’s potential energy. Subsequently, the complex was solvated in a periodic water box using the SPC water model, and sodium and chloride ions were added to maintain a salt concentration of 0.15 M. The complex underwent an NPT (constant pressure, constant temperature) equilibration phase, followed by a 100 ns (nanoseconds) production run in the NPT ensemble. The NPT ensemble was used to simulate systems under constant temperature and pressure, common conditions in biological systems. Trajectories from the simulation were analyzed using Gromacs tools, including root mean square deviation (RMSD), root mean square fluctuation (RMSF), radius of gyration (RG), solvent accessible surface area (SASA), and hydrogen bonding (H-Bond). These analyses provided insights into various structural and dynamic properties of the simulated system, such as shape, flexibility, and interactions with the surrounding solvent.

Additionally, MMPBSA (Molecular Mechanics Poisson-Boltzmann Surface Area) calculations were performed on the ligand-target complex using the last 50 ns of Gromacs trajectories. Complex structures were prepared for calculation in Gromacs, including explicit solvent addition and topology file generation. The g_MMPBSA software was employed to set up the MM-PBSA calculation, and energy decomposition was conducted on each complex over the last 50 ns of the trajectory. This analysis allowed for the determination of binding affinity and the contributions of different energy components to the overall binding energy.

## Results and Discussion

### Investigation of Aromatase Inhibition Potential in Okra Phytocompounds

To elucidate potential aromatase inhibitors within Okra-derived phytocompounds, we conducted a thorough computational analysis of binding affinities. Employing molecular docking techniques, we assessed the binding efficiency of these compounds. The results, summarized in Table 2, provide insights into binding affinity and estimated Ki values for nine Okra compounds in comparison to the commercial aromatase inhibitor, Letrozole.

**Table 2:**
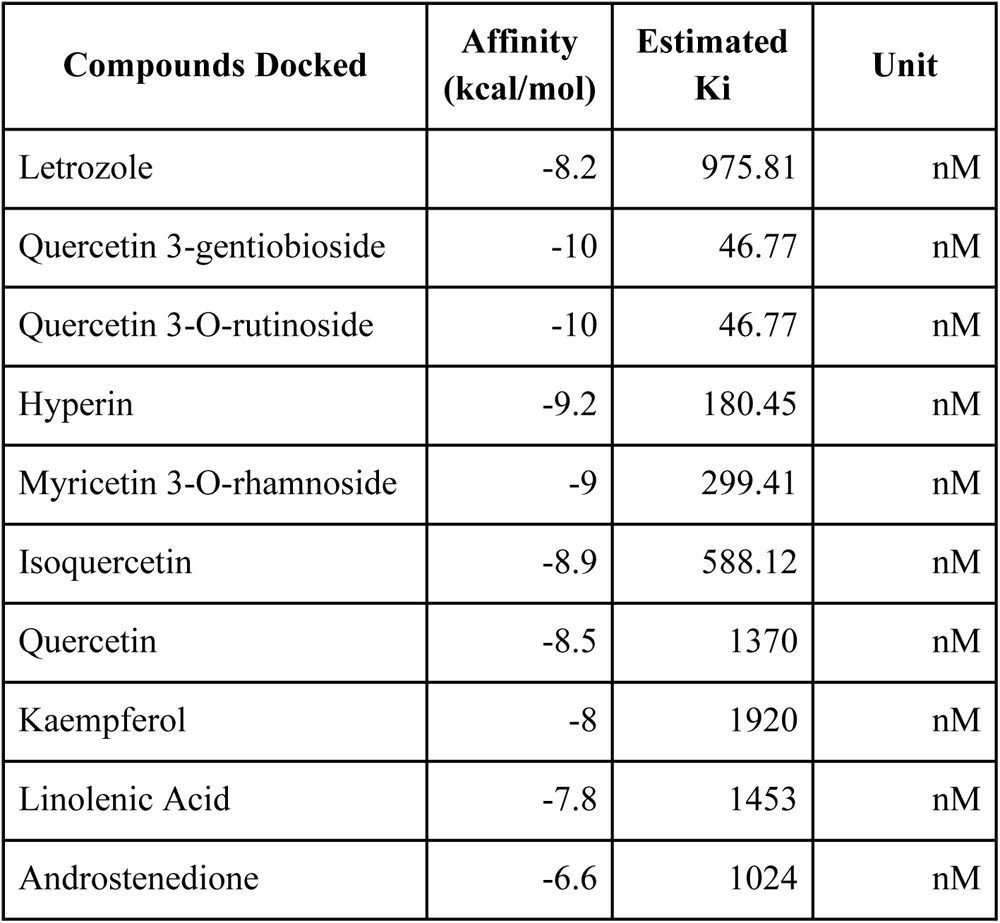
Molecular Interactions of aromatase with Okra phytocompounds.

Notably, Quercetin 3-gentiobioside and Quercetin 3-O-rutinoside exhibited exceptional binding profiles, boasting a remarkable binding affinity of -10 kcal/mol and an estimated Ki of 46.77 nM. Hyperin closely followed suit. Comparing these findings with Letrozole, a third-generation commercial AI with an affinity of -8.2 kcal/mol, underscores the potential of these novel Okra compounds as effective alternatives in the realm of aromatase inhibition.

Furthermore, our analysis suggests a marginally enhanced binding stability for Quercetin 3-gentiobioside and Quercetin 3-O-rutinoside when contrasted with Letrozole. This observation further bolsters the case for considering these Okra-derived compounds as promising alternatives in the field of aromatase inhibition.

### Exploration of Molecular Interactions with Quercetin 3-Gentiobioside

In light of its remarkable binding affinity, we conducted a comprehensive examination to unveil the intricate molecular interactions between Quercetin 3-gentiobioside and the aromatase protein. These interactions play a pivotal role in delineating the effectiveness of binding inhibition. In particular, our focus centered on elucidating hydrogen bond interactions by comparing Quercetin 3-gentiobioside with both the natural substrate, Androstenedione, and the commercial AI, Letrozole. A summary of our findings is presented in Table 3.

**Table 3:** Hydrogen Bond Interactions of Letrozole and Quercetin 3-gentiobioside.

| Compound | Name |
| --- | --- |
| Letrozole | TRP141, ARG435 |
| Androstenedione | MET374 |
| Quercetin 3-gentiobioside | ARG115, ARG145, ARG435, THR310 |

Quercetin 3-gentiobioside exhibited a significantly higher number of hydrogen bond interactions when juxtaposed with the natural substrates, Androstenedione and Letrozole. Impressively, Quercetin 3-gentiobioside engaged in four distinct hydrogen bond interactions with specific residues, namely ARG115, ARG145, ARG435, and THR310. Intriguingly, among these interactions, Arg435 also participated in a hydrogen bond interaction with Letrozole. This observation indicates that Quercetin 3-gentiobioside binds within the same binding pocket as the commercial drug, underscoring its potential as an aromatase inhibitor.

Moreover, we investigated non-bonded interactions for both Letrozole and Quercetin 3-gentiobioside. Letrozole displayed a total of nine interactions, comprising four weak hydrogen bonds and the remaining non-bonded interactions, notably hydrophobic interactions. In contrast, Quercetin 3-gentiobioside exclusively engaged in six weak hydrogen bond interactions, devoid of any hydrophobic interactions. A detailed breakdown of these interactions is provided in Table 4.

**Table 4:** Non-bonded Interactions of Letrozole and Quercetin 3-gentiobioside.

| Compound | Name | Type of Bond |
| --- | --- | --- |
| Letrozole | TRP141 | Hydrogen |
|  | ARG435 | Hydrogen |
|  | CYS437 | Hydrogen |
|  | ALA438 | Hydrogen |
|  | ILE133 | Hydrophobic |
|  | ALA306 | Hydrophobic |
|  | ALA438 | Hydrophobic |
|  | ILE133 | Hydrophobic |
|  | ALA306 | Hydrophobic |
| Quercetin 3-gentiobioside | ARG145 | Hydrogen |
|  | THR310 | Hydrogen |
|  | ARG115 | Hydrogen |
|  | CYS437 | Hydrogen |
|  | ALA306 | Hydrogen |
|  | ALA306 | Hydrophobic |
|  | ALA306 | Hydrophobic |
|  | ALA438 | Hydrophobic |
|  | CYS437 | Hydrophobic |
|  | ILE132 | Hydrophobic |
|  | ILE133 | Hydrophobic |
|  | ALA438 | Hydrophobic |

### Evaluation of Compound Safety and Pharmacological Properties

In order to assess the safety and pharmacological characteristics of the top-performing compounds, namely Quercetin 3-gentiobioside, Quercetin 3-O-rutinoside, Hyperin, and Myricetin 3-O-rhamnoside, we conducted an ADMET analysis. This comprehensive evaluation allowed us to compare these compounds based on various critical parameters.

Among the quartet of compounds scrutinized, Quercetin 3-gentiobioside emerged as particularly noteworthy due to its classification as “Practically Non-toxic” in terms of acute oral toxicity. In contrast, the other three compounds were categorized as slightly toxic, in accordance with the guidelines set forth by the US Office of the Federal Register in 2012.

Furthermore, all four compounds exhibited positive interactions with estrogen and androgen receptors, highlighting their potential for modulating hormonal pathways. Encouragingly, none of these compounds displayed indications of carcinogenicity or the ability to penetrate the blood-brain barrier, bolstering their safety profiles.

Given its distinction as the least toxic among the compounds, Quercetin 3-gentiobioside emerges as a strong candidate for further consideration. The detailed findings from this analysis are summarized in Table 5.

**Table 5:** ADMET Profile.

| Compound | ADMET Predicted Profile | Value | Probability |
| --- | --- | --- | --- |
| Quercetin 3-gentiobioside | Blood Brain Barrier | - | 0.75 |
|  | Carcinogenicity | - | 0.99 |
|  | Hepatotoxicity | + | 0.5429 |
|  | Acute Oral Toxicity | IV | 0.4763 |
|  | Estrogen receptor binding | + | 0.8171 |
|  | Androgen receptor binding | + | 0.6413 |
| Quercetin 3-O-rutinoside | Blood Brain Barrier | - | 0.85 |
|  | Carcinogenicity | - | 1 |
|  | Hepatotoxicity | + | 0.6625 |
|  | Acute Oral Toxicity | III | 0.5971 |
|  | Estrogen receptor binding | + | 0.7901 |
|  | Androgen receptor binding | + | 0.6176 |
| Hyperin | Blood Brain Barrier | - | 0.775 |
|  | Carcinogenicity | - | 1 |
|  | Hepatotoxicity | - | 0.5196 |
|  | Acute Oral Toxicity | III | 0.4045 |
|  | Estrogen receptor binding | + | 0.747 |
|  | Androgen receptor binding | + | 0.7633 |
| Myricetin 3-O-rhamnoside | Blood Brain Barrier | - | 0.85 |
|  | Carcinogenicity | - | 0.99 |
|  | Hepatotoxicity | - | 0.5447 |
|  | Acute Oral Toxicity | III | 0.5184 |
|  | Estrogen receptor binding | + | 0.7235 |
|  | Androgen receptor binding | + | 0.6849 |

### Molecular Dynamics Simulations and Binding Analysis

To gain deeper insights into the interaction dynamics between the inhibitor and the target protein, as well as to explore their effects on protein activity, potential binding sites, and inhibitor mechanisms, we conducted a series of dynamic simulations. This section presents the results of our comprehensive analysis.

#### Simulation Setup

The dynamic simulations involved three key scenarios: (i) Protein in its unbound state [3EQM-APO], (ii) Protein bound to Androstenedione [3EQM-ASD], and (iii) Protein bound to Quercetin 3-gentiobioside [3EQM-QUR].

#### Stability Assessment

We initiated our analysis by examining the Root Mean Square Deviation (RMSD) of the backbone atoms in these protein complexes over a 100 nanosecond (ns) period. The RMSD values for the three complexes, 3EQM-APO, 3EQM-ASD, and 3EQM-QUR, were found to be 0.31+/-0.035 nm, 0.29+/-0.069 nm, and 0.29+/-0.16 nm, respectively. These results indicated slight conformational changes during the simulation, but the deviations did not significantly differ from those observed in the unbound protein. Detailed plots are provided in Figure 3, affirming the relative stability of these protein-compound complexes throughout the simulation. The small differences between the complexes suggest a high degree of structural similarity, likely attributable to their shared structural components.

**Figure 1:**
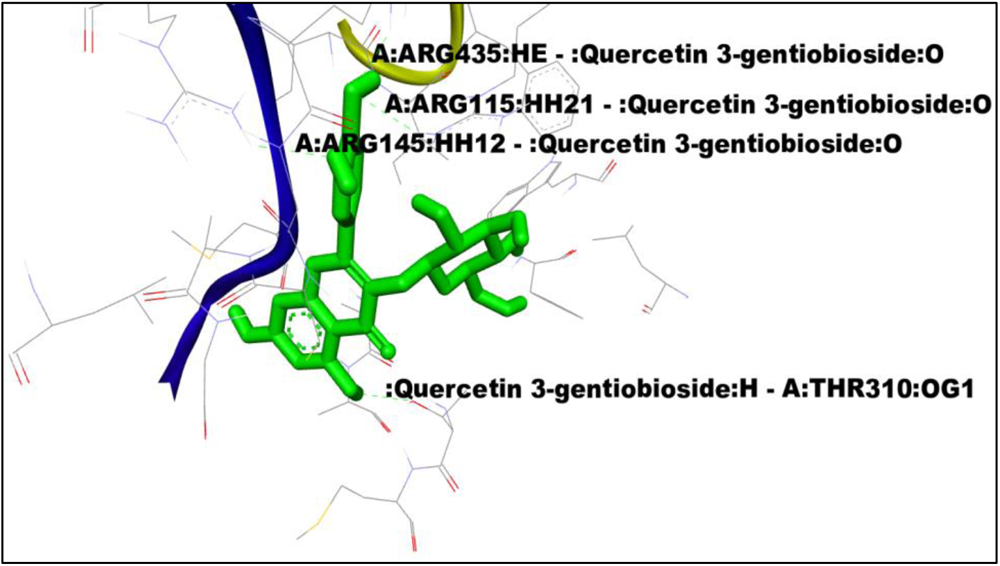
Hydrogen bond interactions of Quercetin 3-gentiobioside.

**Figure 2:**
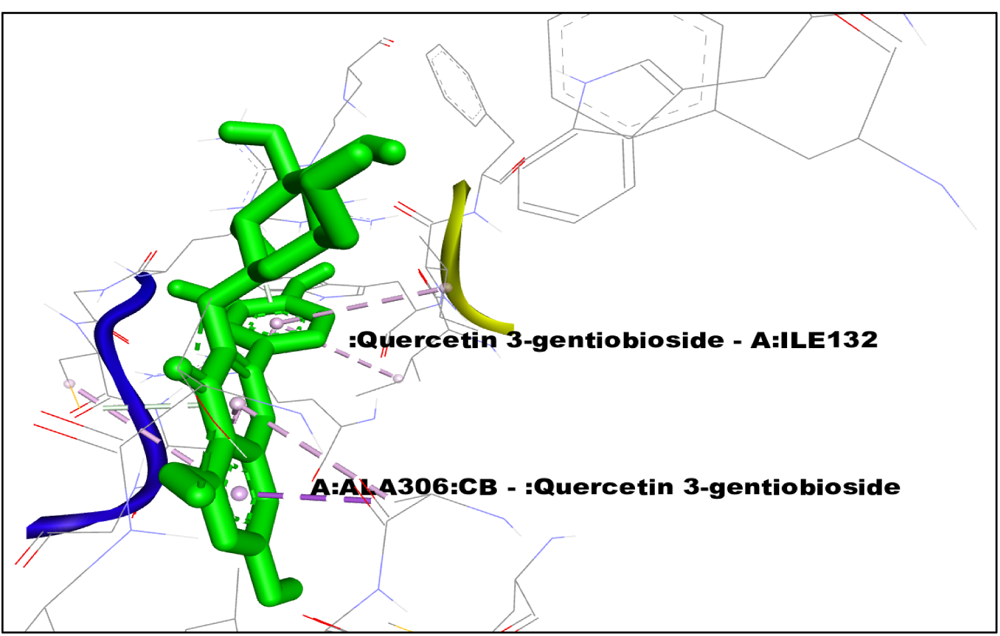
Non-bond interactions of Quercetin 3-gentiobioside.

**Figure 3:**
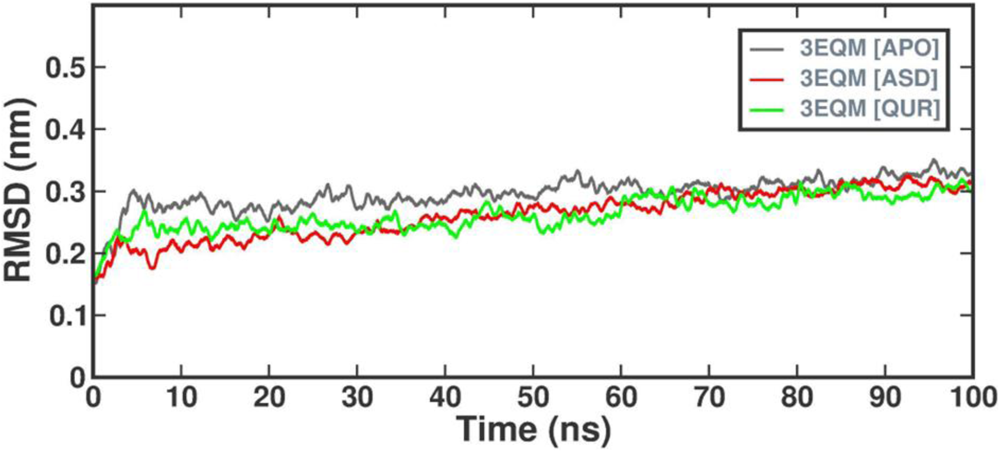
Root Mean Square Deviation of Backbone Atoms Protein Flexibility Analysis.

To assess the impact of ligand presence on protein stability, we conducted Root Mean Square Fluctuation (RMSF) analysis over the 100ns simulation period. RMSF values were determined to monitor vibrations and structural changes. Encouragingly, our results, as illustrated in Figure 4, indicated that no significant structural changes occurred during the simulation.

**Figure 4:**
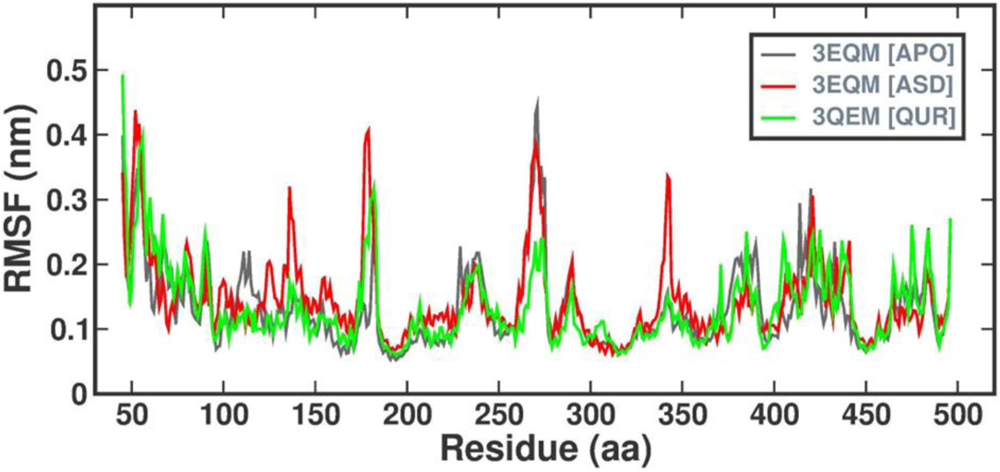
Root Mean Square Fluctuation of C-alpha Atoms.

#### Structural Compactness Evaluation

The Radius of Gyration (Rg) was employed to assess the overall structural compactness of the protein complexes during the simulation. Figure 5 and the accompanying data revealed that 3EQM-APO, 3EQM-ASD, and 3EQM-QUR complexes exhibited a similar pattern of Rg values throughout the simulation, with average Rg values of 2.28+/-0.014 nm, 2.29+/-0.019 nm, and 2.28+/-0.019 nm, respectively. These results suggest that the protein complexes maintained their structural stability over time.

**Figure 5:**
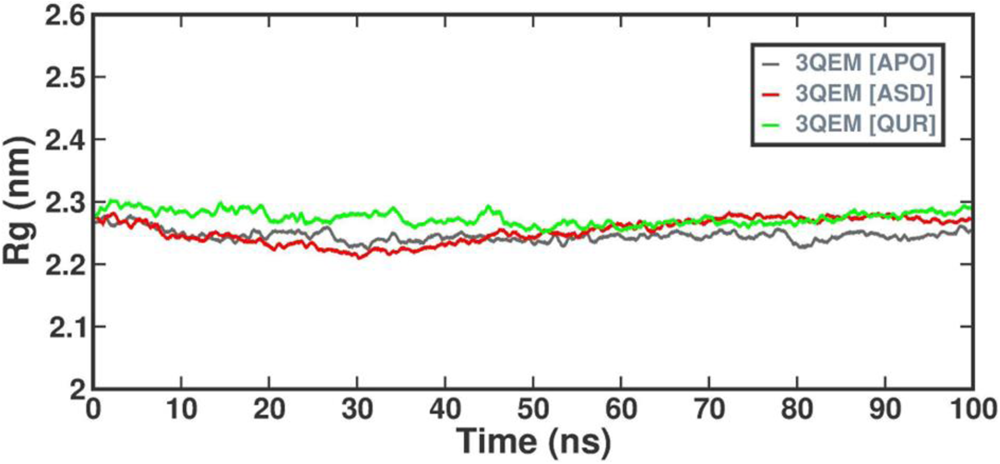
Radius of Gyration (Rg) of Backbone Atoms Hydrophobic Core Analysis.

To investigate the compactness of the hydrophobic core within the protein, we analyzed the change in Solvent Accessible Surface Area (SASA) over time (Figure 6). The average SASA values for 3EQM-APO, 3EQM-ASD, and 3EQM-QUR protein complexes were found to be 210.01+/-4.45 nm, 209.46+/-3.51 nm, and 221.16+/-5.61 nm, respectively. These observations suggest that the structural integrity of the protein remained consistent throughout the simulation.

**Figure 6:**
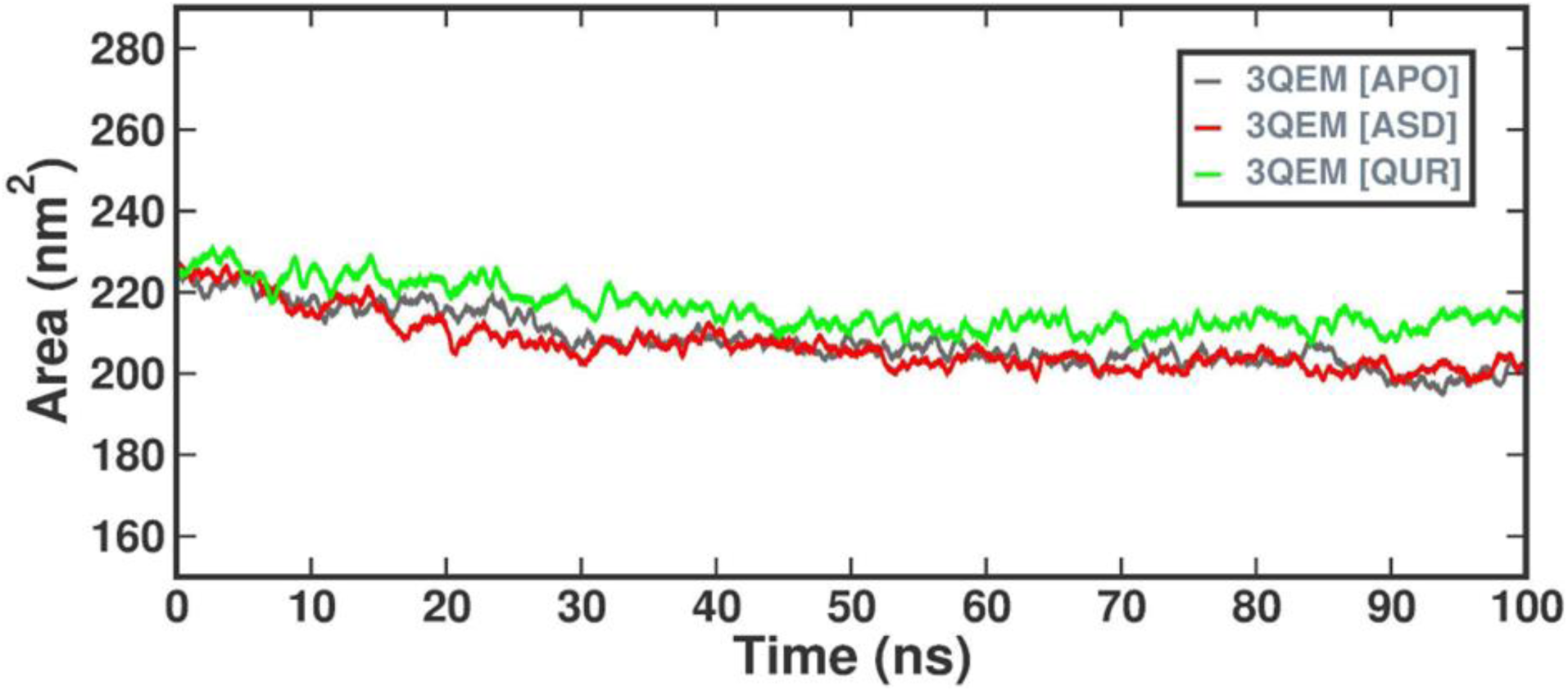
Change in Solvent Accessible Surface Area (SASA) of Backbone Atoms.

#### Hydrogen Bond Analysis

In our pursuit to evaluate the stability of protein-ligand complexes, we confirmed the hydrogen bonds identified during molecular docking through simulation analysis. Figure 7 presents the hydrogen bond interactions within the 3EQM-ASD and 3EQM-QUR complexes.

**Figure 7:**
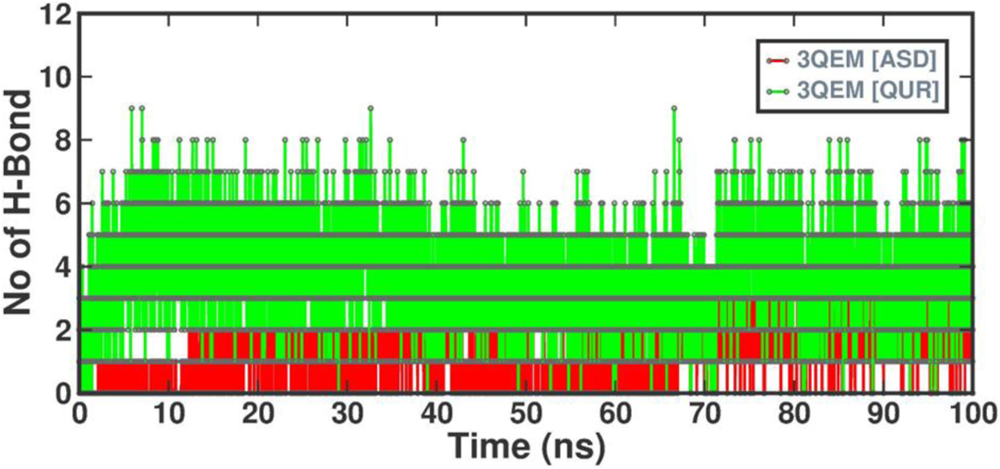
Hydrogen Bond Analysis.

#### Binding Affinity Assessment

To quantify the binding affinity of 3EQM-ASD and 3EQM-QUR, we employed the MM-PBSA method to compute the interaction energy. Table 6 compares the energy contributions of van der Waals, electrostatic, polar solvation, and overall binding energies between these two complexes. Notably, 3EQM-QUR exhibited a van der Waals energy of -216.179 +/- 15.056 kJ/mol, electrostatic energy of -77.298 +/- 15.584 kJ/mol, polar solvation energy of -77.298 +/- 15.584 kJ/mol, and a binding energy of -122.265 +/- 16.461 kJ/mol. These values were compared to those of 3EQM-ASD, the natural substrate of aromatase, shedding light on the efficient and superior binding of 3EQM-QUR.

**Table 6:** Binding Strength Analysis by MM-PBSA Method.

| System | van der Waal energy | Electrostatic energy | Polar solvation energy | Binding energy |
| --- | --- | --- | --- | --- |
| ASD | $-173.365 \pm 8.604$ kJ/mol | $-26.652 \pm 7.125$ kJ/mol | $107.384 \pm 9.412$ kJ/mol | $-109.846 \pm 8.816$ kJ/mol |
| QUR | $-216.179 \pm 15.056$ kJ/mol | $-77.298 \pm 15.584$ kJ/mol | $-77.298 \pm 15.584$ kJ/mol | $-122.265 \pm 16.461$ kJ/mol |

In conclusion, our dynamic simulations and binding analysis provide valuable insights into the interactions and stability of protein-ligand complexes, supporting the potential efficacy of 3EQM-QUR as an aromatase inhibitor. Further investigations are warranted to confirm these findings and advance our understanding of these molecular interactions.

In summary, aromatase inhibitors are a class of drugs used to treat Estrogen positive breast cancer. Commercial AI’s have captured the current market but come with the cost of different side effects like hot flashes and vaginal dryness, to name a few. Plant based AI’s have proven to be a promising alternative. Okra is a popular vegetable with good nutritional significance. There have been several studies done with the bioactive compounds of Okra against a variety of diseases, like diabetes and problems with digestion (Elkhalifa *et al*., 2021) . Okra is a flavonoid rich food source. These flavonoids are known to possess anticancer, anti-inflammatory, and antiviral properties (Kopustinskiene *et al*., 2020) . Nine such flavonoids were chosen for this study and compared to the commercially available AI letrozole. On evaluation, Quercetin 3-gentiobioside showed the best binding affinity among the nine compounds and was found to be better than Letrozole. This compound was further studied for its binding efficacy, where Quercetin 3-gentiobioside showed a similar binding to the protein when compared to letrozole. ADMET analysis for the first four compounds was performed as they had good binding affinity. Out of the four, Quercetin 3-gentiobioside was found to be practically non-toxic, whereas the other three compounds namely Quercetin 3-O-rutinoside, Hyperin, and Myricetin 3-O-rhamnoside were characterized as slightly toxic. Hence, Quercetin 3-gentiobioside was further considered for molecular dynamics. The results of the MD simulation carried out for 100 nanoseconds confirm the stability of the 3EQM-QUR complex, which was contributed by the formation of a significant number of H-bonds and other non-bond interactions. Overall, our study findings suggest that Quercetin 3-gentiobioside may act as an efficient aromatase inhibitor. However, further preclinical and clinical research is needed to confirm the findings of this study.

## Conclusion

This research has contributed valuable insights into the intricate interactions between Aromatase and potential inhibitors derived from okra, as elucidated through molecular docking and dynamic simulations. The rising incidence of breast cancer has driven the exploration of novel phytocompounds sourced from plants like okra, known for their potential anticancer properties. While Aromatase Inhibitors (AIs) have shown promise in the management of breast cancer, improving survival rates, they are not without their associated side effects.

Our study has unveiled the exciting possibility that compounds found in okra could be strong candidates for preclinical evaluation, with Quercetin 3-gentiobioside emerging as a particularly noteworthy contender for the role of an aromatase inhibitor. This discovery hints at a promising future for these okra-derived compounds as potential Aromatase Inhibitors, offering hope for more effective and better-tolerated treatments in the ongoing battle against breast cancer. Further research and clinical trials are warranted to explore their full therapeutic potential, bringing us one step closer to improved breast cancer management.

## Acknowledgement

The authors would like to acknowledge the Sri Ramachandra Institute of Higher Education and Research (DU) for their constant support.

